# Chelation Therapy Using Static Magnetic Field

**DOI:** 10.1101/429597

**Authors:** Jyoti Shaw, Ayan Chakraborty, Sanjoy Chatterjee, Maitreyee Bhattacharyya, Anjan Kr Dasgupta

## Abstract

Static magnetic field (SMF) is reported to mimic chelation agents. In thalassemia blood, SMF minimizes iron overloading effect. In such cells ROS level is reduced, in presence of moderate strength (70mT). Pilot study on transfusion dependent thalassemia, thalassemia carriers and normal red blood clearly indicate the differential effects of SMF on thalassemia, thalassemia carrier and normal cells. SMF also reduces ROS induced DNA damage in lymphocytes. A comparative study on the iron chelating drug deferasirox and SMF treated lymphocytes further confirm our main thesis. This non-invasive therapeutic regime for hemochromatosis may serve the dual purpose of treating, iron overloading and minimizing, cellular damage by lowering the ROS level.

## 1. Introduction

An article published few years back [1] highlighted the dual objectives of chelation therapy. Traditionally it has been used for treating clinical conditions associated with iron overload, typically in transfusion-dependent thalassaemia major. An additional goal of this therapeutic regime, has been to minimize the cellular damage caused by iron accumulation. In this paper we propose a new methodology that enables chelation like effect without any direct addition of chelating agent.

The proof of our concept of this new form of chelation was inspired by one of our earlier work in which we showed the differential effects of static magnetic field on normal and cancer cells [2]. In 2011, Tao and Huang reported changes in blood viscocity in response to static magnetic field [3]. Such scientific reports give useful insights about the sensitivity of physiological systems to magnetic field.

Conservative therapies like using magnetic field to regulate blood pressure was practiced in medieval period [4]. At present, magnetic field is used for diagnostic purpose such as Magnetic Resonance Imaging (MRI), is a non-invasive method, which is able to penetrate from surface to deep tissues with precise and high spatial resolution (10–100nm)[5, 6]. In the last fifteen years, use of magnetic nanoparticles in biomedicine and oncology had been revolutionized for the treatment and diagnostic regimes [7].

Iron congestion is one of the severe consequences in thalassemia patients. It mainly originates from regular blood transfusions, ineffective erythropoiesis and increased gastrointestinal iron absorption which leads to iron overload in the body [8]. As a result, excess iron accumulates in the system which affects the organs and impairs the immune system, placing the patients at greater risk of infection and illness[9]. In normal condition iron always remain bound to a protein such as transferrin or ferritin because free iron is toxic to cells as it generates free radicals from normal metabolites via fenton reactions[10]. In thalassemia, due to transferrin supersaturation, excess iron remains free in the system. These free iron pool is labile and toxic and causes huge damage to the cells by generating free radicals. This condition is commonly referred to as iron overload or haemochromatosis [11].

Commonly, iron chelators are used to chelate free iron present in the system. At present three drugs are mainly available which are given to the patients. These are Desferrioxamine (DFO), Deferasirox (DFX) and Deferiprone (DFP). Among these chelation drugs, only Desferrioxamine and Deferasirox are approved by FDA. However, these iron chelator drugs have some side effects which vary from patient to patients[12]. Thus there is a requirement to remove excess iron from cellular environments with iron chelation therapy (ICT) in order to avoid the serious clinical sequelae associated with iron overload in patients. But due to such negative side effects imparted by these iron chelating drugs, ICT is still a major reason for unsatisfactory compliance. Therefore, long term use of these chelation drugs is not suitable and is also very expensive for a patient. Therefore, it is very essential to develop an alternative therapy which overcomes these shortfalls of the conventional drug chelation therapies. In the present study, we had shown the chelation effects of static magnetic fields on thalassemic patients.

## 2. Materials and Method

### 2.1 Selection and evaluation of the study subjects

The protocol was duly approved by the ethical committee of Medical College (MC/KOL/IEC/NONSPON/325/09-2014), Calcutta (Kolkata). Samples were collected after obtaining written informed consent from the subjects. The selection and evaluation of study subjects are detailed in our recent publication[13]. In the present study, blood sample were collected from the subjects in EDTA and heparin vacutainer, for RBC and lymphocyte isolation, respectively with informed consents as prescribes by the ethical committee.

### 2.2 Oxidative stress assay in red blood cells

100µl of the EDTA blood collected from the subjects were processed to prepare packed red blood cells. Firstly, blood was diluted with 3ml of 0.89 percent saline and then centrifuged at 1200 rpm for 5 min. This step was repeated twice in order to remove all white blood cells. Then, RBC pellet was washed with phophate buffered saline (PBS). The RBCs were suspended in PBS at a concentration of one million RBCs per ml. Then RBC suspension was divided in four sets. One set was incubated with static magnetic (100mT) for one hour at room temperature. In another set, iron chelating drug, deferasirox (DFX) was added at 100uM concentration. In the third set, DFX was added and incubated in presence of SMF of same strength. The fourth set was control. After incubation, oxidative stress or ROS level was determined using probe DCFDA [14] and run in BD influx flow cytometer (from Becton Dickinson). The FCS-format data was analyzed using FlowJo LLC.

### 2.3 Ex vivo treatment of lymphocytes

Firstly lymphocytes were isolated and treated from both thalassemia patients and controls following the protocol described by Shaw et al [13]. Lymphocytes suspension were first divided into two sets of tubes. Each set contained four tubes and subsequent treatment protocol was similar as described in the referred publication. One set was incubated in presence of SMF (100mT) and another set was kept as control. Each experiment was done in triplicate. Then oxidative stress and DNA damage assay were performed for such treated cells.

### 2.4 Oxidative stress in lymphocytes of thal patients

For the measurement of oxidative stress in lymphocytes and RBCs, probe 2,7-Dichlorofluoresceindiacetate better known as DCFDA was used. DCFDA is a retroactive fluorescent probe that enters a live cell where the diacetate moeity is cleaved by the cellular esterase. In presence of ROS, DCF is oxidized to give the active fluorescent dye. The sample to be assayed was incubated with 10µM for 15–20 min in the dark. Then sample was washed and resuspended in PBS. The sample was analyzed immediately in the spectrofluorimeter (PTI,Quantamaster TM40) having excitation wavelength at 485nm and emission was scanned around 525nm.

### 2.5 DNA damage analysis by Comet Assay

After treatment, a part of lymphocytes were tested for the presence of DNA damage by Comet Assay. The detail procedure was described in detail in Shaw et al, 2014. The slides were then observed under microscope and at least 100 cells were scored for each slide. Triple slides were prepared for each sample. The comets were scored in the COMETSCORE program.

## 3. Results

### 3.1 Differntial clustering of RBCs from different samples

In fig 1, the RBCs from different samples form different cluster in a forward scatter versus side scatter density plot. These patterns were found to be similar within a group like thalassemia group or normal group. In flow cytometry, the position of population clusters in a morphological plot is the characteristics of a particular cell type. Thus such distribution of RBC clusters may be used as a signature for a particular group. RBCs distribution for thalassemia and control group was different. Whereas, RBCs from thalassemia carriers formed clusters similar to both control group and thalassemia major group.

**Fig 1.**
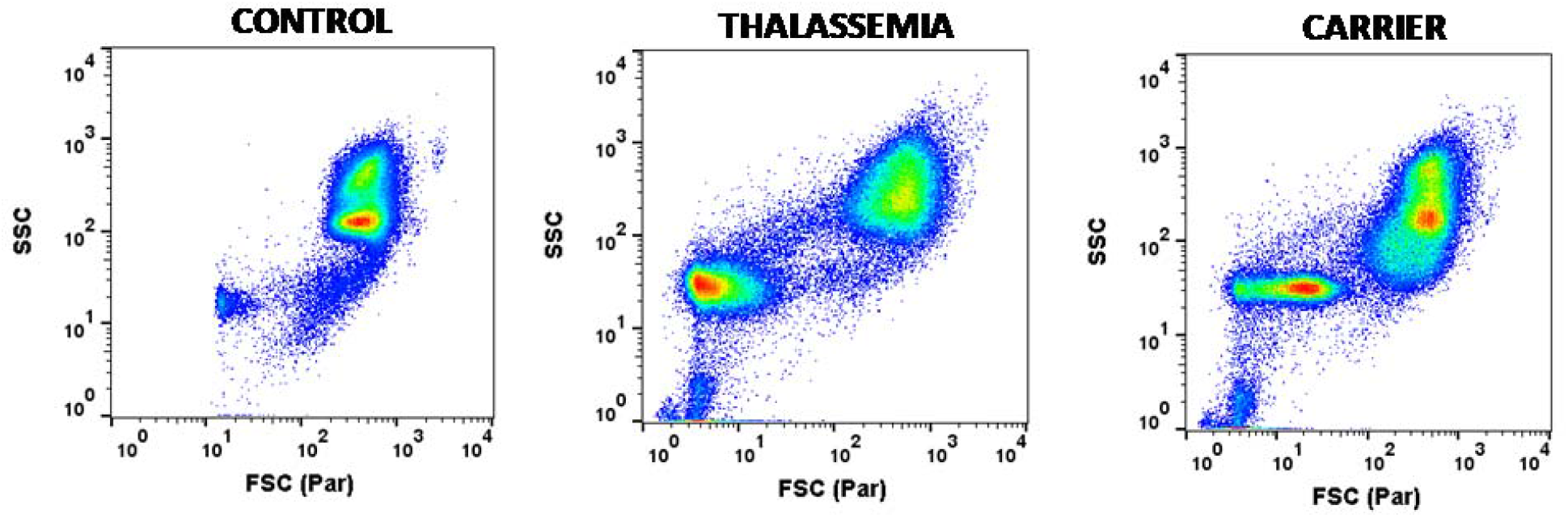
Differential clusters of thalassemic RBCs. The figure shows the psuedocolor density plot of side scatter versus forward scatter in three different human samples. Left panel graph represent control sample, middle graph represent thalassemic samples and right graph sample from thalassemic carrier.

### 3.2 Alteration in RBCs distribution in response to iron chelators

In fig 2, the effect of iron chelator drug Deferasirox (100µM) was tested for the three samples of control, thalassemia and thalassemic carrier. Adjunct histograms was also displayed alongside of the scatter plots. Blue dots in the scatter plot represent untreated and red dots are cells treated with drug. Two graphs are superimposed to understand the change in distribution of cells. No change in pattern was observed for control RBCs but shift in clusters was observed in case of both thalassemia and carriers. More RBCs shifted towards the right in response to the drug.

**Fig 2.**
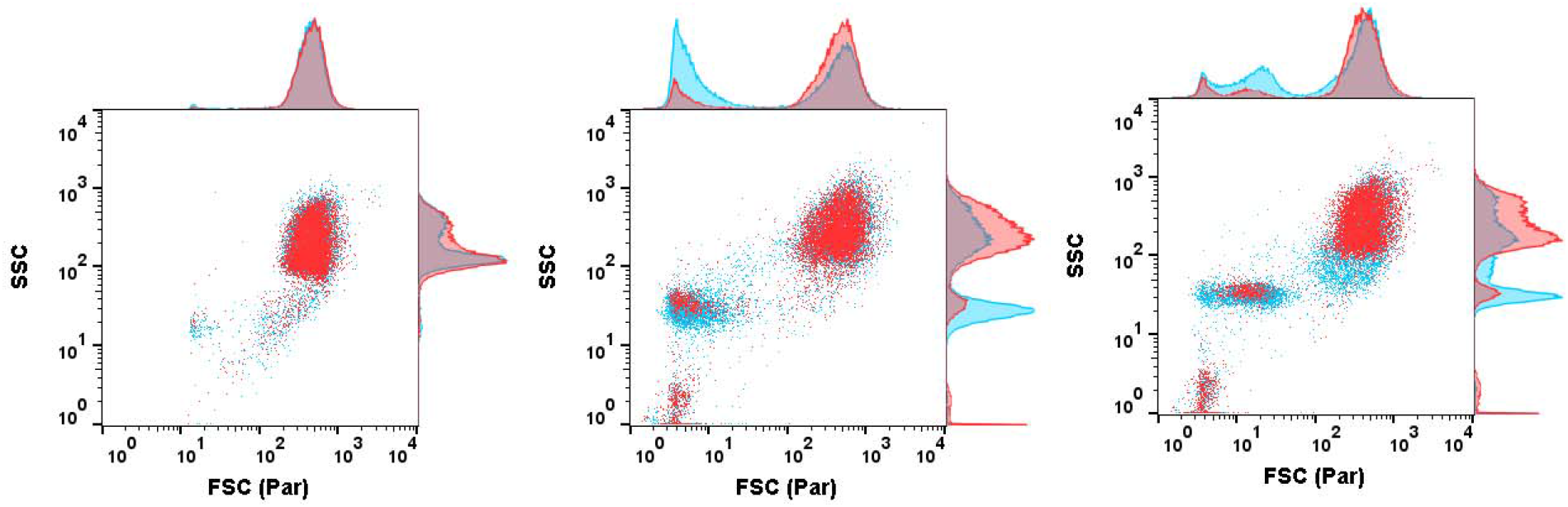
Effect of Iron chelators on RBC population. The figure shows the dot plot of RBCs from three types of samples. Data of the same sample at two different condition are superimposed to understand the change in population distribution. Blue color dots represent untreated sample whereas red color dots represent sample treated with iron chelator DFX (100µM). Left panel shows data from control sample, middle panel is of thalassemic sample and right panel is of thalassemic carrier

### 3.3 Alteration in RBCs’ distibution in response to SMF

Similar effect was observed in case of cells exposed to SMF as shown in fig 3. Like fig 2, same nomenclature was followed in the fig 3. In this case also, control RBCs had no effect due to magnetic field as the population clusters completely overlap in the superimposed graphs but both thalassemia and carriers cells responded to SMF. Due to SMF treatment, RBC clusters having lower value of forward scatter completely shift towards right as well as there is also increase in side scatter. Keen observations of fig 3 revealed that there was more upward shift in side scatter of carrier cells as compared to thalassemia majors.

**Fig 3.**
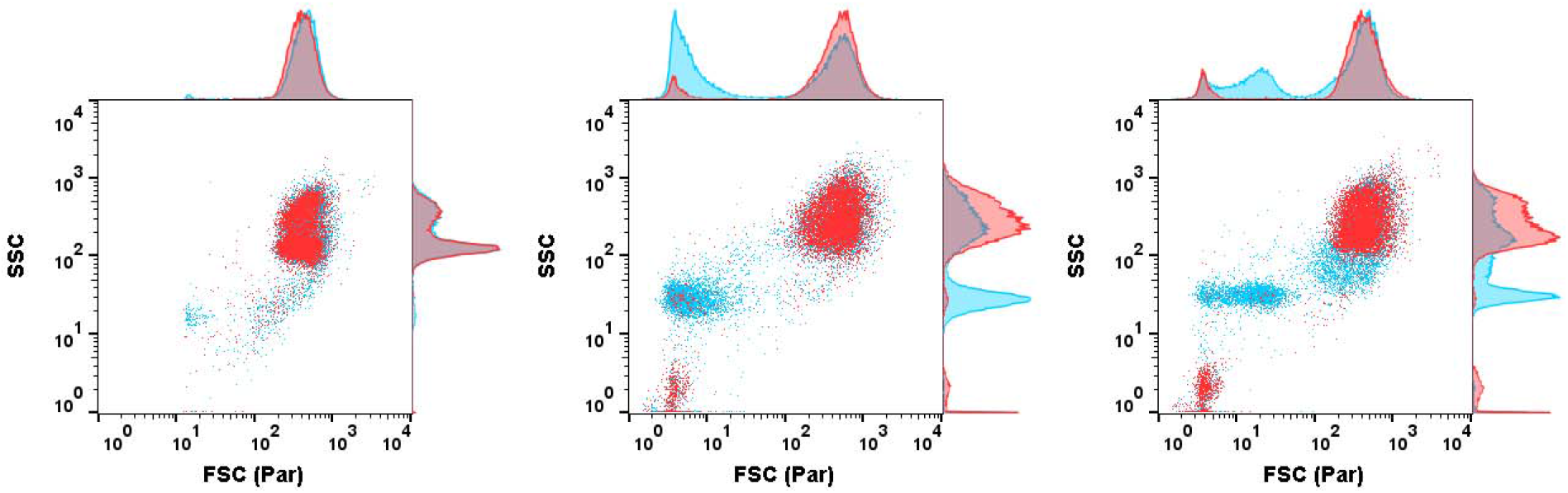
Sensitivity of RBCs to static magnetic field (SMF). The figure shows the dot plot of RBCs from three types of samples. Data of the same sample at two different conditions are superimposed to understand the change in population distribution. Blue color dots represent untreated sample whereas red color dots represent sample incubated in presence of SMF. Left panel shows data from control sample, middle panel is of thalassemic sample and right panel is of thalassemic carrier

Thus it shows that there is a group of RBC cells present in the diseased condition but absent in control or healthy individuals which respond to the drug and SMF.

### 3.4 Reduction in ROS level in RBCs treated with Deferasirox and or SMF

The above treatments (namely Deferasirox and SMF) not only altered morphological properties of cells but also affected the functional properties of RBCs. It is observed that endogenous ROS level decreased in all the three groups of cells namely control, thalassemia patients and carriers. The results are tabulated below in table 1.

**Table I:** Chelation effect of SMF on thalassemic RBCs

|  |  | -SMF | +SMF |
| --- | --- | --- | --- |
| **Endogenous ROS** | Control | 22.29 | 17.22 |
|  | Patient | 92.15 | 56.5 |
|  | Carrier | 94.3 | 43.6 |
| **Deferasirox** | Control | 36.6 |  |
|  | Patient | 57.7 | 56.7 |
|  | Carrier | 63.6 | 73.8 |

Both Deferasirox and SMF reduced the endogenous ROS level in thalassemia patients and carriers significantly. The values shown in the table represent mean value for a particular group. In case of control RBCs, endogenous ROS level was significantly less as compared to other two groups. SMF further reduced the ROS level in control RBCs but Deferasirox had reverse effect on the same cells. Deferasirox increased ROS level in control samples. The synergistic effect of both iron chelator and SMF was observed when the RBCs of thalassemia were treated with both DFX as well as exposed to SMF. The ROS level in such sets were lesser as compared to DFX alone. In case of carriers variable results were obtained. The mean ROS level in carriers was slightly more as compared to patient group which reduced in presence of SMF or deferasirox but synergistic effect of both SMF and deferasirox was not observed in carrier RBCs.

### 3.5 SMF effect on ROS level in lymphocytes of thalassemia patients

Similar effects of SMF and iron chelator Deferasirox was observed in thalassemic lymphocytes (fig 4). SMF reduced the endogenous ROS level significantly (p=0.0036) as compared to unexposed lymphocytes of thalassemic patients. It also significantly controlled the ROS generation due to external iron salts (p=0.0104) in lymphocytes treated with iron salt (ferrous sulfate). Reduction in ROS level was maximum when both SMF and DFX (iron chelator) was present but the difference between SMF exposed and unexposed ROS level in deferasirox treated lymphocytes was not significant. Similarly, ROS level in lymphocytes treated with both iron salt and chelator in presence of SMF was low as compared to absence of SMF. These results showed the synergism between SMF and iron chelator in controlling ROS generation in lymphocytes of thalassemic patients.

**Fig 4.**
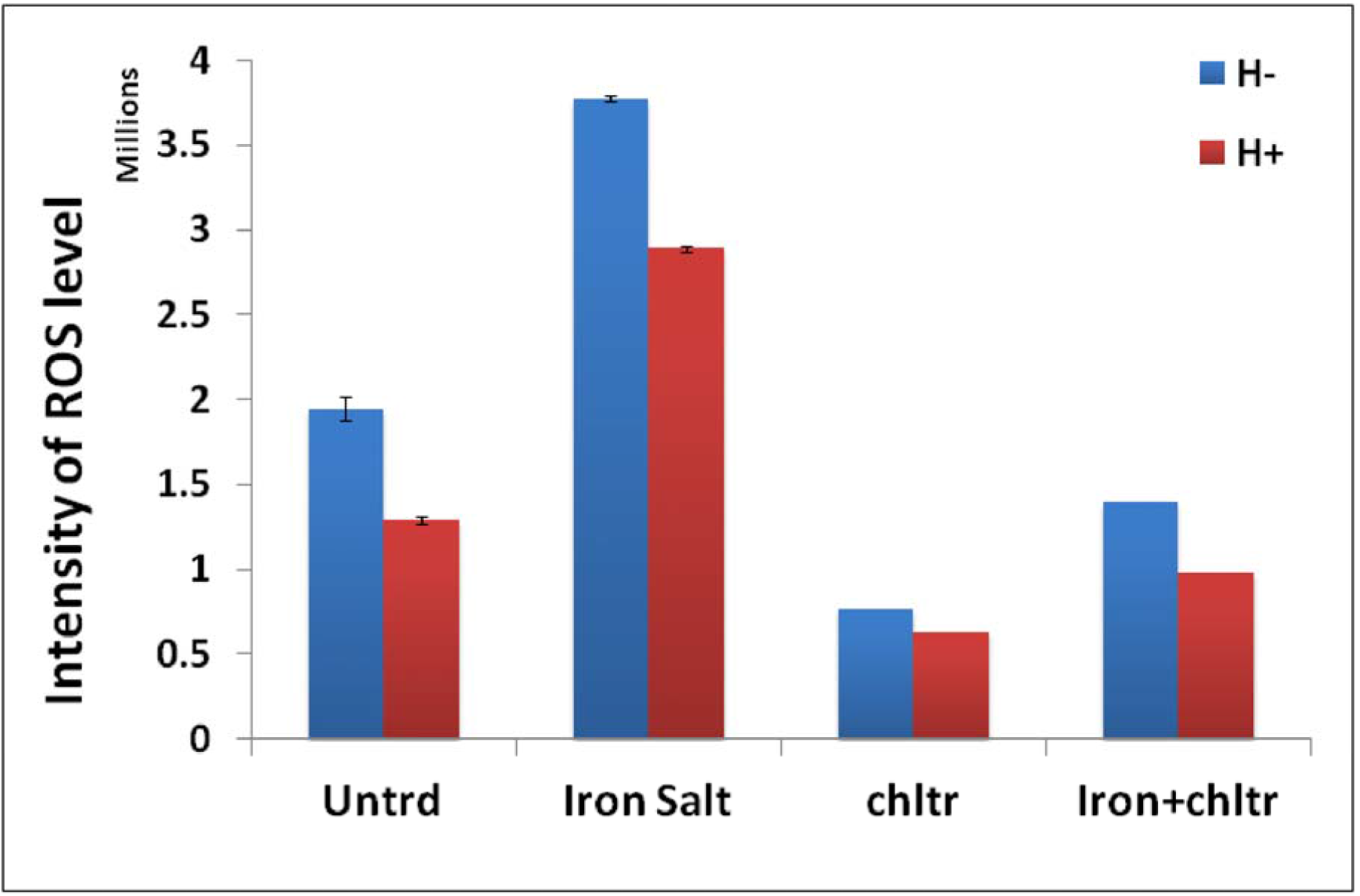
Diminution in intracellular ROS level in lymphocytes in presence (H+) and absence (H-) of SMF in thalassemic lymphocytes. The bar-graph shows the four conditions such as ‘untr’ for untreated, ‘iron salt’ for ferrous sulfate treated cells, ‘chltr’ for iron chelator DFX and ‘iron+chltr’ for cells treated with both iron salt and chelator (details in section 2.3). The results proved the chelation effects of SMF

### 3.6 SMF prevents the increase in ROS in lymphocytes for longer period

SMF was found to reduce endogenous ROS in lymphocytes during prolong incubation till five hours (fig 5). Iron chelator DFX, also controlled ROS generation during prolong incubation as compared to SMF. The synergistic effect of DFX and SMF was again observed in this case (fig 5). SMF significantly controls inflation in ROS level in lymphocytes in absence of any chemical agent during prolong incubation *in vitro*. Deferasirox also controls ROS generation in lymphocytes but efficiency of DFX is more in presence of SMF. Thus synergestic effect of SMF and DFX is also present here.

**Fig 5.**
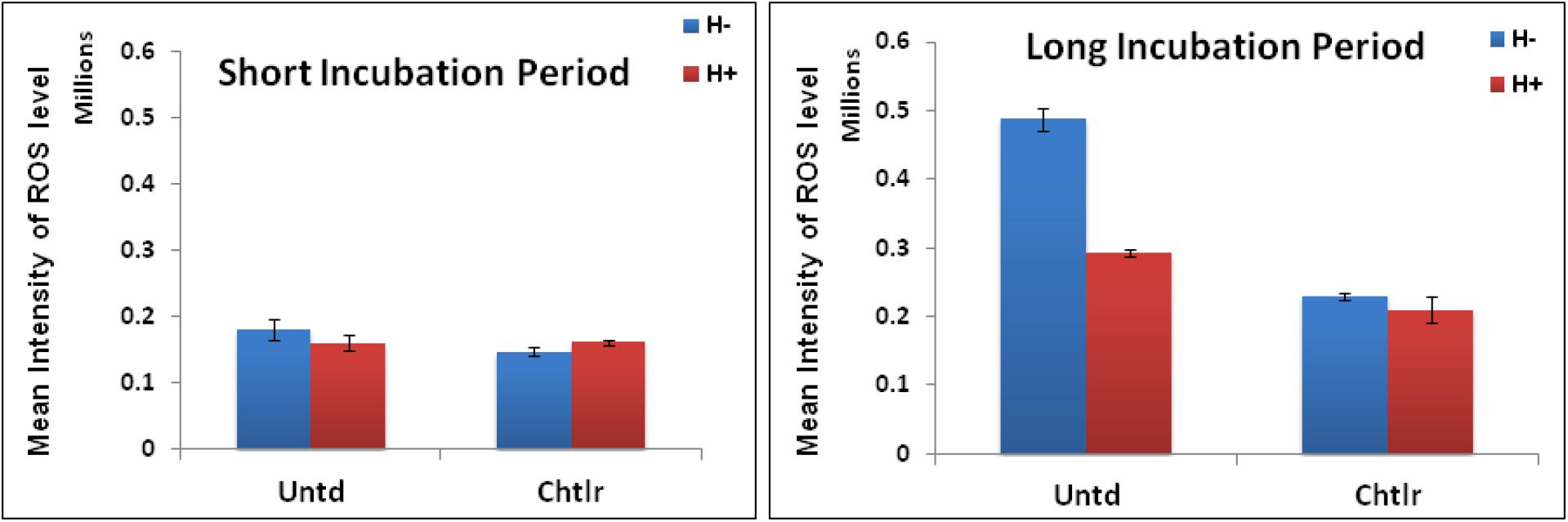
Effect of Incubation period in ROS level in thalassemic lymphocytes in presence (H+) and absence *(H-) of SMF*. The bar-graph shows result of untreated cells as ‘untrd’ and cells treated with DFX as ‘chltr’ Panel A shows result of short incubation period and panel B shows result of long incubation period. During long incubation, synergistic effect of SMF and DFX was observed

### 3.7 SMF reduced endogenous DNA damage in lymphocytes of thalassemia patients

SMF was found to reduce endogenous DNA damage in lymphocytes of thalassemia patients when incubated in presence of SMF *in vitro* (Fig 6). The fig 6, shows result of two such thalassemic cases. Iron chelator, Deferasirox also reduced DNA damage but much less as compared to SMF alone. Lymphocytes treated with Deferasirox and also incubated in presence of SMF, had very less DNA damage as compared to Deferasirox alone. This observation was true for most of the thalassemic cases in the study population. SMF significantly reduced DNA damage in lymphocytes in all the conditions in this experiment. The results of fig 6, shows lymphocytes incubated in presence of SMF alone had least DNA damage as compared to lymphocytes treated with Deferasirox and incubated in presence of SMF. The endogenous ROS level in both carriers and control was found to be negligible.

**Fig 6.**
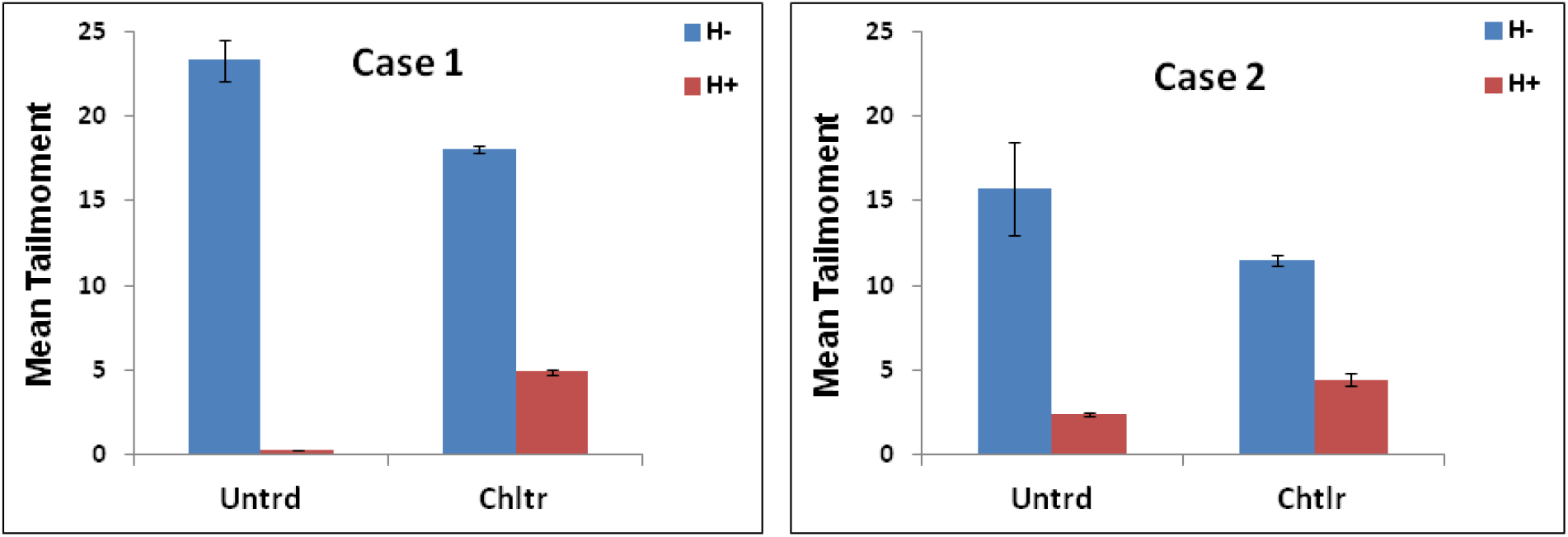
Diminution of endogenous DNA damage of thalassemic lymphocytes in presence (H+) and absence (H−) The bar-graph shows result of comet assay performed in untreated lymphocytes (untrd) and lymphocytes treated with DFX (chltr). The two panel shows results of two such thalassemic cases

## 4. Discussion

The present study vindicates the hypothesis in Shaw et al [2] where we suggested that normal cells are somehow insensitive to magnetic effect. In that paper our compraison group consisted of normal and cancer cells. In the present paper we compare normal cells with thalassemic cells. We find again that magnetic field is unable to perturb normal healthy cells whereas thalassemic cells change their ROS status upon application of magnetic field. Like magnetic field data, normal cells was again unaffected by approved iron chelation agents like Deferasirox (see figure 2 and 3).

The important aspect is that both chelating agents and magnetic field lowers the ROS level and this is apparant in flow cytometry results as well as direct spectroscopic measurements. RBCs of the three type of samples (control, thalassemia carrier and thalassemia major) show a remarkably distinguished pattern of FSC-SSC plot. Such pattern of clusters may be used as a representative for a particular type of cells (say for thalassemia). The distinctive pattern of carrier may be given a special note as this may be a simple way of classifying carrier on a large scale basis.

Now, two population of cells are clearly visible in samples of thalassemia patients and carriers (figure 1) whereas only one cluster is visible for control. Addition of either magnetic field or drug seems to cause a population drift in case of patients and carriers. It appears that in this bimodal population of RBCs in diseased cells or carriers, the smaller sized cluster additionally present was responsive to either magnetic field or chelating agents. The population drift was more pronounced in case of magnetic field (as compared to the chelator) suggesting the future possibility of use of field for treating the iron overloading or haemochromatosis.

The ratio of population levels in two clusters, will provide us the simple quantitative estimate of iron induced damaged cells. On the other hand, the efficient merging of the population in post treatment condition (i.e post magnetic field or chelation agent added conditions) would lead to a metric expressing the responsiveness of the treatment type to the chelation or SMF application. We thus have both diagnostics and therapy which are essence of theranostics. The non-invasive nature of the magnetic field induced theranostic effect deserves a special mention.

The highlights of the present work are:

- presence of two FSC-SSC clusters in thalassmic cases and those from thalassemic carriers
- presence of single FSC-SSC cluster in normal cells
- responsiveness of the lower FSC clsuter to static magnetic field and chelating agent
- apparent reduction of ROS level by static magnetic field implying close relation of ROS level with iron overloading
- chelation like effects induced by static magnetic field, unfolding possibility of noninvasive theranostic treatment of iron overloadding.

The two may not be unrelated as any unpaired spin and SMF would cause significant changes in the chemical activities. For example a triplate oxygen state may behave as singlet in presence of SMF. Similar arguments may hold good for ferric-ferous redox pair. Perhaps SMF induced perturbation of the ferous-ferric redox couple towards the ferrous state may be the key for such chelation behaviour. After all removal of ferric ion is the primary goal of the chelation therapy. This novel non-invasive mode of chelation therapy.

## Compliance with Ethical Standards

***Funding Agency**: Indian Council of Medical Research(ICMR). Project file no. 5/3/8/322/2016.Title: Differential synchronous-A new paradigm for probing oxidative stress and inflammation using serum samples*.

***Conflict of Interest**: There is no conflict of interest in the work*.

***Ethical approval**: All procedures performed in studies involving human participants were in accordance with the ethical standards of the ethical committee of Medical College (MC/KOL/IEC/NON-SPON/325/09-2014)*

